# Spatiotemporal patterns in Golden-cheeked Warbler breeding habitat quality and quantity

**DOI:** 10.1101/2021.03.25.436967

**Authors:** Lindsay M. Dreiss, Paul Sanchez-Navarro, Bryan Bird

## Abstract

The Golden-cheeked Warbler, *Setophaga chrysoparia*, is a migratory songbird listed as endangered under the federal Endangered Species Act that breeds exclusively in central Texas and is heavily impacted by habitat conversion. The species relies on mixed Ashe-juniper and oak woodlands for nest-building and shelter during spring and early summer months. Using land cover data spanning the last 25 years, we conduct a geospatial analysis to quantify changes and\ identify shifts in breeding habitat quantity and quality. Since 1985, 13% of all forests within the warbler’s breeding range were disturbed, with greater incidences near San Antonio (32%) and Austin (24%) metropolitan areas. Additionally, data show a 45% decrease in high-quality habitat (i.e., intact mixed or evergreen core forests) and a decrease in patch size. Habitats within protected areas see a less sharp decline in habitat quality and large increases in warbler sightings, but these only represent 10% of all highest-quality habitat in the breeding range. Drastic declines in habitat quality suggest that generalized metrics of conversion may underestimate true habitat loss as degradation may impact the ecological viability of remaining forests for warbler nesting. Further evidence suggests that the few protected areas within the Texas range continue to play a significant role in warbler breeding. This information will assist researchers and managers prioritizing conservation action and will inform upcoming species status determinations.

---

Habitat loss and degradation through anthropogenic landscape modification are major drivers of declining global wildlife populations and serve as primary threats justifying species’ listing under the United States Endangered Species Act (ESA; Hanski 2011, Bairlein 2016, Thompson et al. 2016, Horváth et al. 2019, Leu et al. 2019). Habitat disturbances are even more notable for Neotropical migrant songbirds that travel long and costly distances between breeding and non-breeding sites. The Breeding Bird Survey of North America – an active roadside census – shows that half of migratory bird species are declining; declines in long-distance Neotropical migrants are more pronounced than those of birds migrating shorter distances (see North American Breeding Bird Survey). These species depend on multiple habitats at different points in space and time, and the reduction in the quality of one habitat can have far-reaching consequences for overall species persistence (Robbins et al. 1989, Zitske et al. 2011, Taylor and Stutchbury 2015, Jackson et al. 2019). Breeding season, though a small proportion of the annual cycles for many migratory birds, is significant due to the more direct association with recruitment and fitness of a species (La Sorte et al. 2017). Therefore, loss and fragmentation of breeding habitats in the United States and other northern locales may have particularly severe ecological implications for imperiled migratory bird populations, especially when breeding distribution is very restricted.

The Golden-cheeked Warbler (*Setophaga chrysoparia*, GCWA) is a migratory songbird listed as endangered under the federal ESA that is heavily impacted by forest conversion. The species is a classic habitat specialist, breeding exclusively in the Edwards Plateau of central Texas, commonly referred to as the Texas Hill Country, and preferring mature, mixed Ashe-juniper and oak woodlands (Pulich 1976, Long et al. 2016). These tree species provide critical material for nest-building and shelter for main GCWA food sources. A wealth of literature assessing the influence of habitat factors on measures of GCWA survival (i.e., presence, reproductive success, nest survival, and density) finds that forest composition, age, and patch size are important to species success (Pulich 1976, Shaw and Atkinson 1990, U.S. FWS 1992, Jetté et al. 1998, Magness et al. 2006, Diamond 2007, Colón et al. 2019). Their specialized preference for this already range-restricted forest type, as well as the high rates of habitat loss from urban developments and transportation infrastructure, led to the listing of GCWA as endangered by the U.S. Fish and Wildlife Service in 1990. Since listing, the threat of habitat loss persists with an estimated 29% reduction in total GCWA breeding habitat between 2000-2010 (Duarte et al. 2013). As of 2020, a very small portion (1.5%) of Texas lands are protected or managed in a way that is consistent with biodiversity conservation (GAP 1 or GAP 2, U.S. Geological Survey 2019). As such, the threat of further habitat loss and degradation remains, and continued monitoring and evaluation of landscapes and populations is necessary to understand the progress of GCWA recovery (Eichenwald et al. 2020).

A mandate of the ESA (section 4(c)(2)), regular reviews of the best available science and commercial information are conducted to revisit population trends, threats to recovery, and accuracy of the listing. Science that demonstrates a range-wide understanding of available breeding habitat conditions and distribution is critical to engaging federal action for proper protections for species recovery (La Sorte et al. 2015). Though coarse, this information may be used to estimate population size and viability and, in the case of habitat conversion, can also serve to evaluate the status and trends of major threats to species recovery (McGowan et al. 2017). For GCWA, analyses are generally focused on habitat quantity, but additional nuances of habitat quality may result in a more refined understanding of population dynamics and can help managers prioritize habitats for conservation and restoration. Most importantly, regular updates to species habitat quality, quantity, location, and use are necessary to understand longer-term temporal trends for better informed conservation efforts and for consideration in regular federal assessments revisiting species listing (and other) decisions. As such, up-to-date spatiotemporal trends in both the quality and quantity of species habitat at larger ecological scales are important for informing conservation management and supporting continued protections for threatened and endangered species.

The last review for GCWA was in 2014 and since then, there have been questions regarding species recovery and listing status. Currently available science on GCWA breeding habitat quantity assesses temporal trends on short timeframes (no more than a decade) and is now a decade out-of-date (Duarte et al. 2013). Additionally, available analyses of habitat quality are restricted to fractions of the breeding range (Loomis Austin 2008, Heger and Hayes 2013). There is a need for an amended breeding habitat assessment to reflect recent landscape changes and data availability as well as to inform upcoming species status assessments and the next steps in conservation planning.

The objective of this study is to conduct a more comprehensive spatiotemporal model of GCWA breeding habitat distribution. As part of this study, we use geospatial data spanning the last 25 years to 1) update range-wide dynamics in GCWA habitat quantity, 2) analyze spatiotemporal patterns in habitat quality, and 3) compare findings with GCWA sightings and with local protected areas for considerations of habitat use and conservation, respectively.

## METHODS

This study focuses on spatiotemporal changes to habitat loss and degradation for the entirety of the Golden-cheeked Warbler (*Setophaga chrysoparia*, GCWA) breeding range. Breeding and nesting activities are confined to central Texas, USA where ideal habitat varies in density and cover. Generally, habitat is more common in the southern and eastern regions of the range.

Nesting habitat is generally defined by the tree species composition. Warblers nest in habitat made up of mature Ashe-juniper and a combination of other species such as live oak (*Quercus fusiformis*), Shallow-lobed oak (*Quercus breviloba)*, Texas oak (*Q. buckleyi*), post oak (*Q. stellata*), blackjack oak (*Q. marilandica*), Lacey oak (*Q. glaucoides*), shin oak (*Q. sinuata*), sugarberry (*Celtis laevigata*), Texas ash (*Fraxinus texensis*), Nuttall’s oak (*Quercus taxana)*, cedar elm (*Ulmus crassifolia*), escarpment cherry (*Prunus serotina var. eximia*), pecan (*Carya illinoinensis*), and little walnut (*Juglans microcarpa*) (U.S. FWS 1992). Quality habitat generally occurs in forest patches at least 100 hectares in size with moderate to high density of older trees. Forests with greater variation in tree height, greater average tree height, and greater density of deciduous oaks are also associated with higher densities of GCWA (Wahl et al. 1990).

### Data Acquisition

All spatial data inputs are publicly available and analyzes focus on locations inside the GCWA range as defined by the U.S. Geological Survey’s Gap Analysis Program (Table 1). Data were acquired in the summer of 2020 and analyses use ArcPro v 2.3 (Esri, USA). Final outputs are available at DOI 10.17605/OSF.IO/T4DJX.

**Table 1.** Data acquired for spatiotemporal analyses on habitat disturbance, quality, and sightings.

| Data | Source | Temporal | Spatial | Analysis |
| --- | --- | --- | --- | --- |
|  |  | Resolution | Resolution |  |
| Landscape change | Eichenwald et al. 2020 | 1985-2018 | 30m | Habitat disturbance |
| Forest Change | MRLC National Land<br>Cover Database | 1985-2016 | 30m | Habitat disturbance |
| Modeled historic<br>land use | Sohl et al. 2018 | 1985 | 250m | Habitat quality |
| NLCD | MRLC National Land<br>Cover Database | 2016 | 30m | Habitat quality |
| Warbler sightings | iNaturalist & eBird | 1934-2020 | point | Sighting Hotspots |
| Protected Areas | U.S. Geological Survey | 2019 | vector | All |
| Urban areas | U.S. Census | 2019 | vector | Habitat disturbance |
| Breeding range | U.S. Geological Survey<br>GAP | 2001 | vector | Study area |

### Spatial Analyses

#### Habitat Disturbance

Google Earth Engine implementation of the LandTrendr algorithm (Kennedy et al. 2018) identify loss of habitat within the GCWA breeding range between 1985 and 2018 from Landsat imagery via breakpoints in temporal trends of NDVI (see Eichenwald et al. 2020). For our purposes, habitat loss is the area where one habitat is degraded quickly over a short period of time (including from natural or prescribed burns). For each year, we calculate the area of disturbed and undisturbed habitat throughout the entire breeding range, in urban and non-urban portions of the range, and in the metropolitan areas of Austin and San Antonio separately as defined by U.S. Census urban area boundaries. We conduct identical calculations with forest loss data from the National Land Cover Database to corroborate the analysis.

#### Habitat Quality

We apply a previously developed habitat assessment framework to determine location and acreage of quality GCWA habitat for the entire breeding range in 1985 and the most current year of the National Land Cover Dataset (2016, Heger and Hayes 2013). Coarser-resolution (250m) models of historical land use and land cover for the contiguous U.S. estimate habitat in 1985 (Sohl et al. 2018). To account for differences in data resolution, we resample and mask the land cover data from 1985 by historic forest disturbance data (NLCD 2016), assuming that all pixels labeled as either a) never experiencing a disturbance or b) experiencing a disturbance after 1985 were forested in 1985. The framework for scoring habitat suitability is based on a large body of literature citing forest composition, landscape fragmentation, and edge effects as related to Dreiss 8

GCWA presence, survival and breeding success (e.g., Pulich 1976, U.S. FWS 1992, Jetté et al. 1998, Magness et al. 2006, Peak 2007, Long et al. 2016, Reidy et al. 2017, Reidy et al. 2018, Colón et al. 2019). Additionally, Heger and Hayes tested multiple models and confirmed that an evergreen and mixed forest-based model performed better than models using only evergreen or mixed forest types. Habitats of highest quality are intact mixed or evergreen forest cores. Factors and criteria used for scoring habitat quality include:

Forest type: where mixed or evergreen forest types and deciduous forest within 100m of mixed/evergreen forest received a 1. All other land cover types received a 0.

Landscape context: neighborhood statistics were determine the percent of forest land cover in a 210m radius. Areas that are 80-100% forested receive the highest score (4) and areas 0-20% forested the lowest (0).

Edge effect: scores are docked 1 point if they are within 100m of the forest edge.

For each dataset, we calculate the amount of habitat by score throughout the entire breeding range. Descriptive statistics are also generated to compare results in urban areas and specifically for Austin and San Antonio metropolitan areas.

#### Hotspot Analysis

Occurrence data from open-source community science databases (iNaturalist and eBird) help assess hotspots in GCWA sightings. Point locations are grouped by date: sightings prior to 1995 (n = 647) are considered more closely linked to historic landscape patterns and sightings after 2010 (n = 14,568) may give more insight on current spatial patterns. Sightings during breeding Dreiss 9 season (April to August) are used. Additionally, only non-duplicate points representing live observations are used if they are associated with an observation date and a meaningful latitude and longitude (not the centroid of the state or county). Kernel density is used to calculate density of GCWA sighting per square kilometer, with the top quartile of density values representing ‘hotspots’ for GCWA nesting. Centrality and directional distribution of sightings are also compared between the two time periods. We have high confidence in drawing conclusions about breeding location based on sighting location because a warbler’s range is on average a 100-m radius around its nest, depending on the quality of habitat (Reidy et al. 2018).

#### Protected Areas Overlay

Community science data and results from habitat quantity and quality analyses are used in overlays to calculate descriptive statistics based on other landscape designations and coverages from the protected areas database of the U.S. (PADUS v 2.0). U.S. Geological Survey’s Gap Analysis Program (GAP) codes are specific to the management intent to conserve biodiversity. GAP 1 and 2 areas are managed in ways typically consistent with conservation and are considered ‘protected’ in this context.

## RESULTS

Before 1985, 25.64% of Golden-cheeked Warbler (GCWA) breeding range was covered by forest lands (over 4.54 million acres). Between 1985 and 2016, 13% of all forests within the warbler’s breeding range were converted to other land uses (Fig. 1). Forest conversion was more extreme in parts of the range in metropolitan areas, with 24% forest loss in the Austin metropolitan area and 32% loss in the San Antonio metropolitan area. Generally, for all regions, there are greater rates of decline in more recent years. Habitat quality also declined during this time period throughout the breeding range (Fig. 2). In the 1980s, over one-tenth of the forested habitat within GCWA breeding range was intact mixed or evergreen core forests. In 2016, high quality habitat made up 5% of the breeding range, indicating a 45% decrease in the highest quality breeding habitat (Fig. 3). Remaining quality habitat is more fragmented, with significantly smaller patch sizes than in 1985 (Fig. 4; *t* = 1.96, *p value* <0.001). Generally, quality habitat is more concentrated along the southeastern extent of the breeding range and some forested areas to the northwest of Austin and in the northern parts of the breeding range have improved since 1985 (Fig. 2). Habitats within protected areas (i.e., GAP Code 1 & 2) see less sharp declines in habitat quality from 27% to 20% of the breeding range (27% decrease; Fig. 3). However, protected habitats currently represent only 10% of all highest-quality habitat in the breeding range.

**Figure 1.**
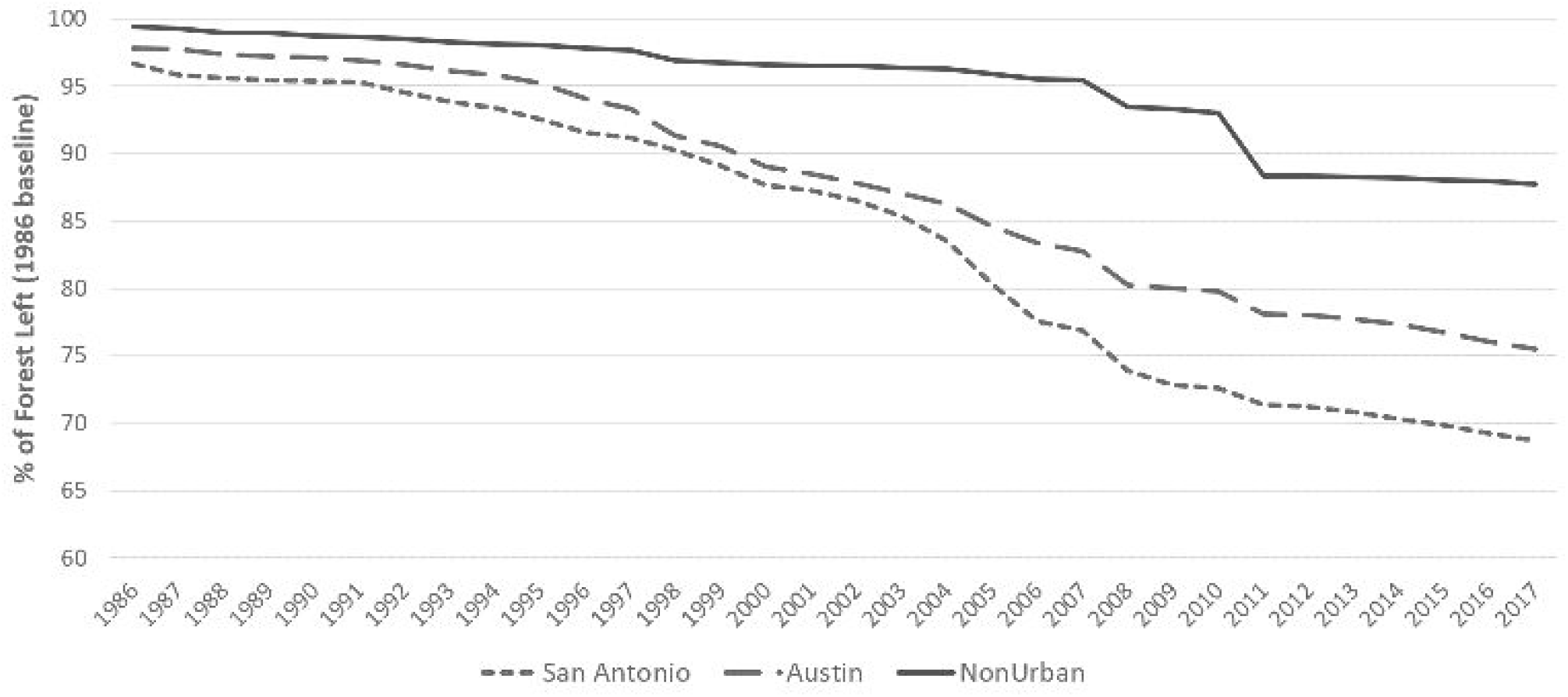
Annual declines in forested land cover relative to a 1985 baseline for portions of the Golden-cheeked Warbler breeding range that fall outside of urban boundaries and portions that fall within the urban boundaries of San Antonio and Austin, respectively.

**Figure 2.**
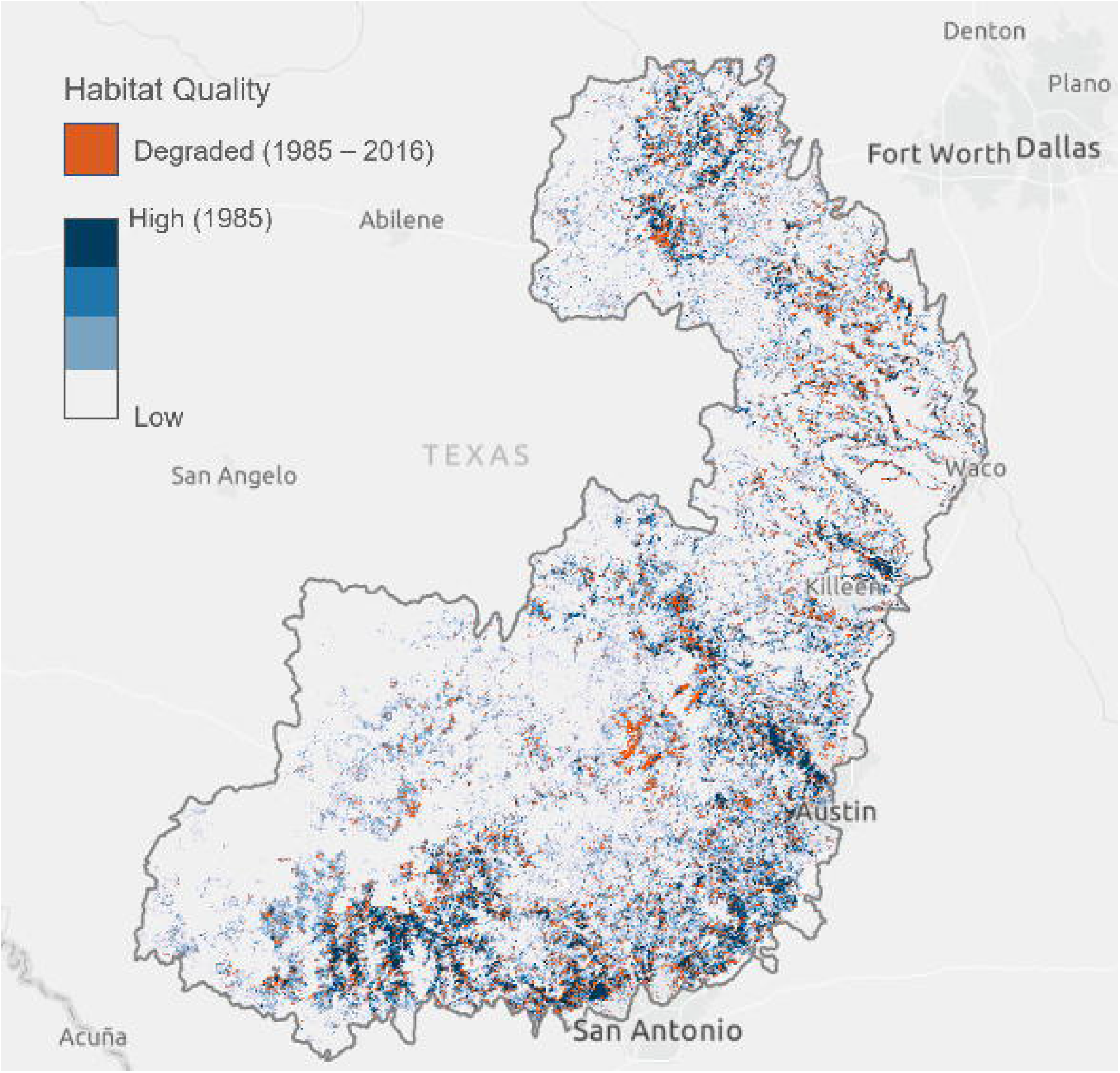
A map of overall change in Golden-cheeked Warbler breeding habitat quality between 1985 and 2016. Blues indicate areas of lower-quality habitat in 1985, white indicates high-quality habitat in 1985, and red indicates high-quality habitat areas that experienced a decline in habitat quality between 1985 and 2016.

**Figure 3.**
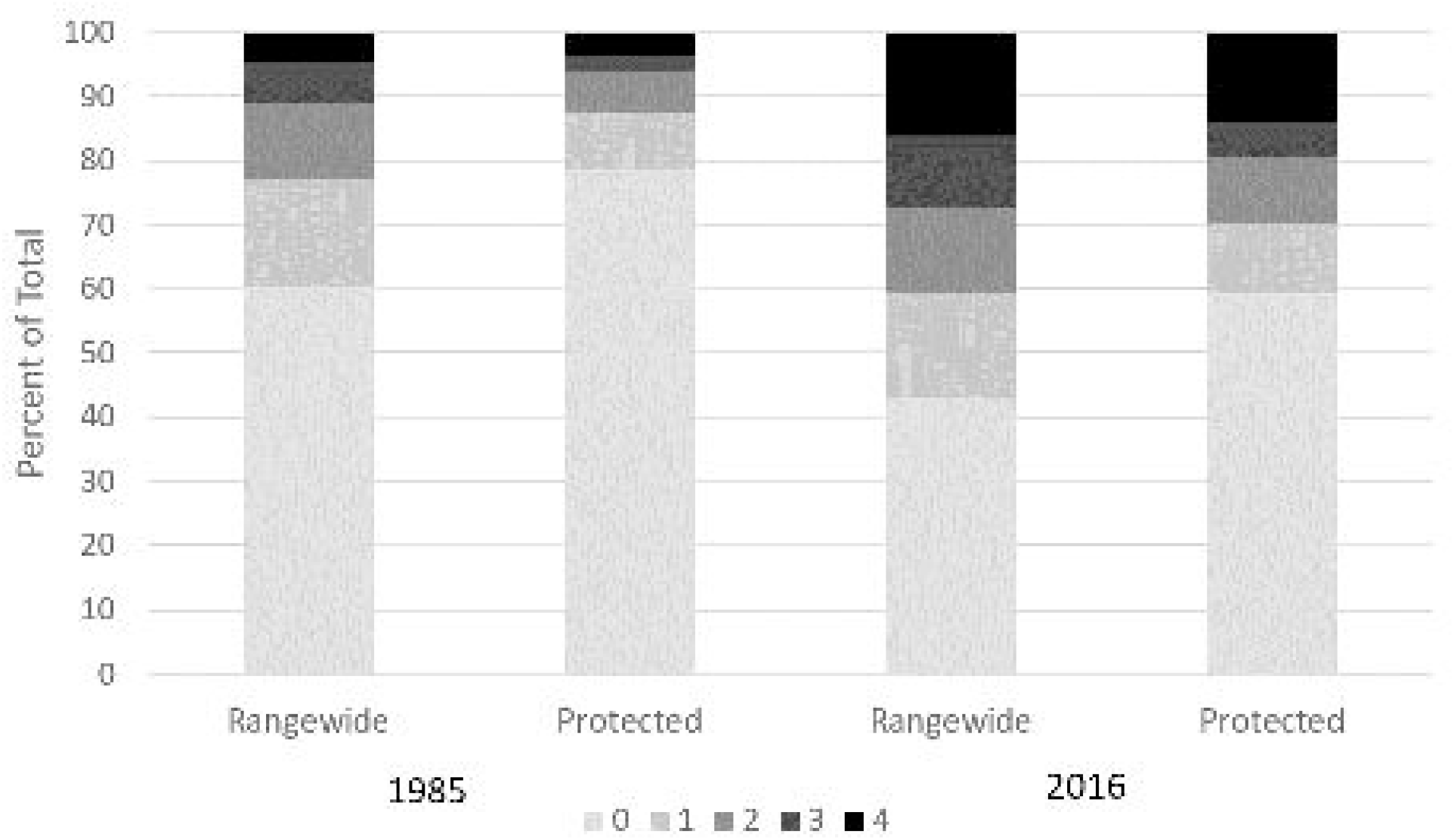
Percent of the Golden-cheeked Warbler breeding range by habitat quality value (0 = low quality, 4 = high quality) for habitats throughout the entire range and for habitats that fall inside protected areas managed for biodiversity conservation (U.S. Geological Survey’s Protected Areas Database of the U.S., GAP codes 1 and 2) for 1985 and 2016, respectively.

**Figure 4.**
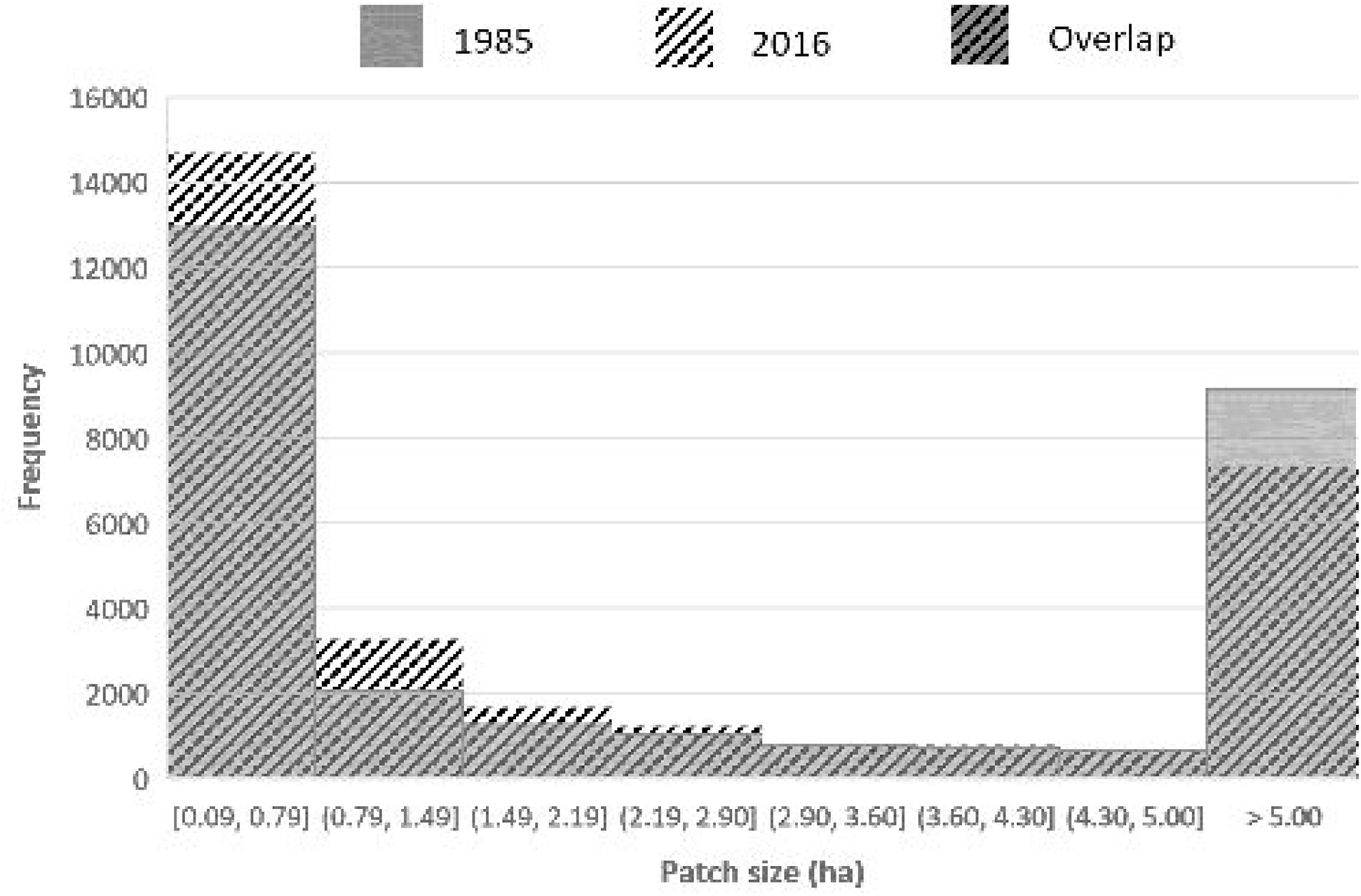
Histograms showing frequency of patch size for high quality habitat areas in 1985 (gray) and 2016 (black hatching) indicate habitat fragmentation over the time period. Means of the two groups were significantly different at α = 0.05 (p value <0.001).

GCWA sightings are generally spatially coincident with habitat quality. In the 1980s, 39% of sightings were in high-quality habitat. As of 2020, sightings in high-quality habitat had dropped to 28%, but this is still disproportionately high given that only 5% of the breeding range consists of high-quality breeding habitat. The proportion of sightings in protected areas in the breeding range has increased dramatically from 5% of sightings before 1995 to 59% after 2010. There were small, localized shifts in the location of GCWA sighting hotspots between the 1980s and 2020, but breeding range-wide, the distribution of hotspots (centrality and dispersion) remains the same (Fig. 4). Sightings were once very concentrated to the southeastern portions of the range, but are now less concentrated, with a few hotspots formed in western parts of the range. Parks in and around San Antonio metropolitan area and Lost Maples Natural Area continue to be hotspots for sightings. All but one of the hotspots were associated with a protected area (Fig. 5).

**Figure 5.**
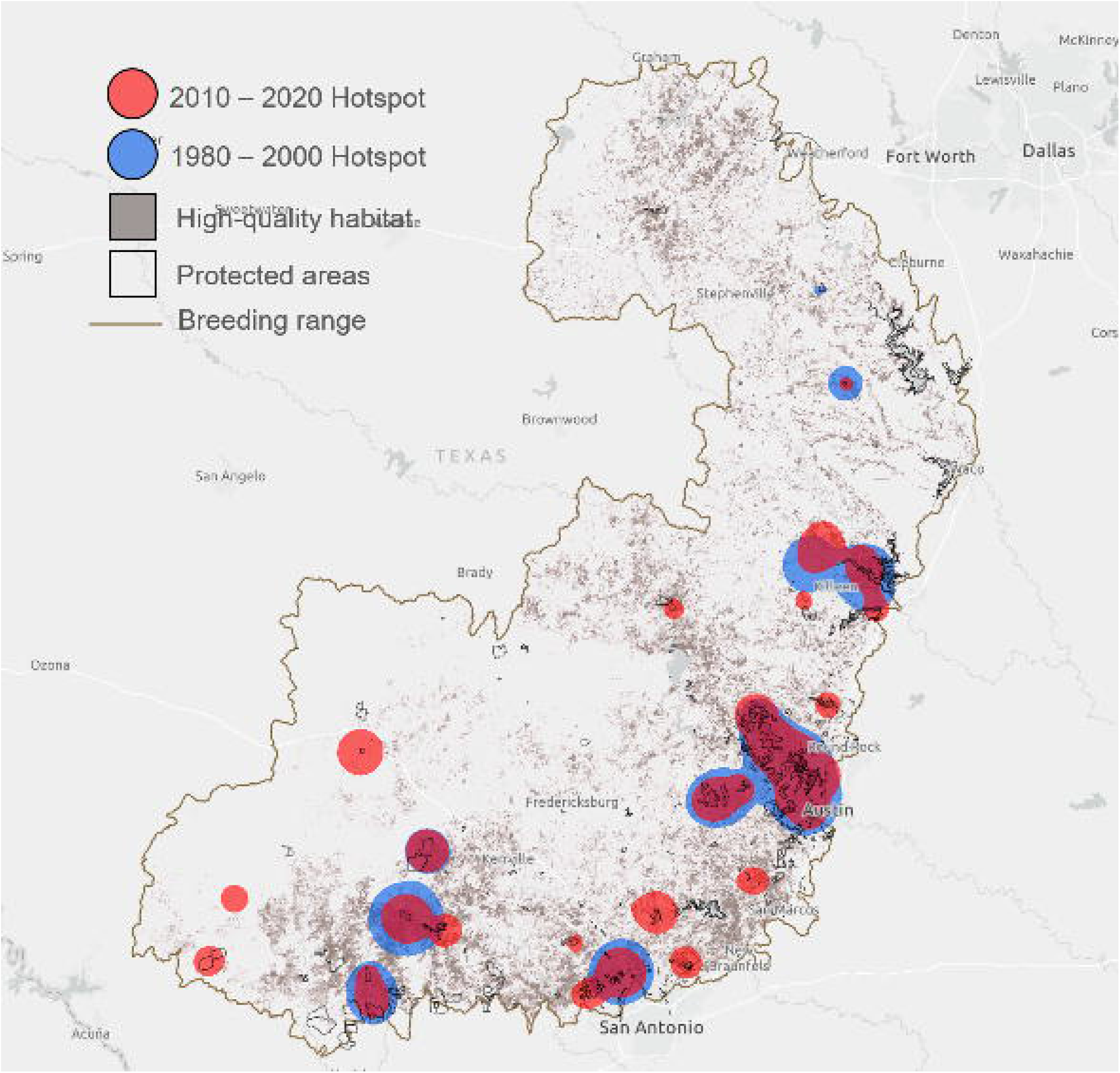
A map comparing locations of high-quality habitat in 2016, protected areas managed for biodiversity conservation (U.S. Geological Survey’s Protected Areas Database of the U.S., GAP codes 1 and 2), and hotspots for Golden-cheeked Warbler sightings between 1980 – 2000 (in blue) and 2010 – 2020 (in red).

## DISCUSSION

We quantify the absolute change in forest cover within Golden-cheeked Warbler (GCWA) breeding range and changes in breeding habitat quality over a 30-year period. Overall, we estimate a 13% loss in breeding habitat with high spatial heterogeneity in landscape conversion closely tied to human developments. This value is lower than other estimates and may be reflective of changing forest dynamics that occur over the longer-term study period-this would support our findings that greater decline has occurred in more recent years (Groce et al. 2010, Duarte et al. 2013). Additionally, drastic declines in habitat quality suggest that 13% is an underestimation of effective habitat loss as degradation may impact the ecological viability of remaining forests for GCWA nesting. The amount of intact core forest habitat fell 45% in the 30-year period leaving quality breeding sites concentrated along the southeastern extent of the breeding range and in protected areas.

Human impact is on the rise in Texas landscapes and may compromise habitat quality. In recent years, Texas has had the largest increases in population of any state in the U.S. (U.S. Census Bureau 2020). In the 30-year period that was studied, the population grew 73.9% and growth is projected to continue at a steady rate to 2050 (88.3% increase in the next three decades; Texas Demographic Center 2019). Within GCWA breeding range, at least four counties are projected to see population increases of over 100% by 2050, all of which coincide with areas of quality habitat in the southeastern parts of the range: Williamson, Hayes, Comal, and Kendall counties.

Increased development pressures in the Texas Hill Country could continue to drive the trends of GCWA habitat disturbance and degradation. Our data indicate that quality forests have undergone fragmentation resulting in smaller habitat patches. Similar trends have been reported more specifically for Ashe-juniper distributions across the state due to an increase in pastureland and development (Diamond 1997). A reduction in canopy cover can lead to decreased nest success for forest songbirds (Martin and Roper 1988, Trzcinski et al. 1999, Twedt et al. 2001). Canopy cover is also essential to conceal GCWA nests located in the mid-story to upper canopies of trees, thus reducing the probability of nest predation and parasitism (Reidy et al. 2008). Additionally, fragmentation of breeding habitat may represent barriers to dispersal of birds and important genetic material (Lindsay et al. 2008). Hence, there is already evidence of notable genetic differentiation among populations of GCWA, having important implications for management of species like GCWA that are relatively vagile, but highly specialized in their habitat preferences. Restoration and protection of connected patches may be the best option for conserving or recovering such species (Young and Clarke 2000).

We found that only 10% of the highest quality forest habitat are in protected areas, creating both challenges and opportunities. These lands, because they are managed in ways consistent with biodiversity conservation, generally represent higher quality habitats with fewer human disturbances (Rosa and Malcom 2020). Our findings indicate that protected areas within GCWA breeding range also exhibited declines in quality, but degradation was buffered relative to the overall range. As human populations grow and landscape conversion continues, protected areas are expected to grow in importance. Nearly all (17 out of 18) of GCWA sighting hotspots from our analysis were associated with a protected area. Additionally, the proportion of sightings that occurred on protected areas saw a significant increase. It should be noted that public lands may Dreiss 13 have a higher proportion of sighting simply due to their accessibility to observers. However, a preliminary analysis of GCWA occupancy models from Morrison et al. (2010) also demonstrates the importance of protected areas to GCWA success: areas with at least 70% probability of occupancy make up 13% of the breeding range, but 62% of protected areas. Public protected areas can play a central role in habitat conservation efforts because they are more amenable to the application of broad-scale management strategies that more closely align with species conservation. However, the extent to which public protected areas can benefit migratory bird populations depends on how well protected areas are represented within the breeding range (La Sorte et al. 2015). Currently, areas managed for conservation (GAP status 1 and 2) represent 3.23% of the breeding range. Lands with more intermediate mandates (GAP 3) provide a higher degree of flexibility for the implementation of management recommendations more closely aligned with maintaining biodiversity. However, these lands are also limited in the state of Texas (1.71%). Expansion of protections to key habitats would require that resources be spent in agency land acquisition or in private lands conservation.

Our findings demonstrate a need for strengthening current conservation measures and expanding upon protections for GCWA habitat to ensure greater breeding success and, ultimately, species recovery. Newer proposals to protect at least 30% of U.S. lands and waters by 2030 to address the biodiversity and climate crises may provide additional opportunities for land designations and conservation efforts for imperiled species like GCWA (Exec Order No 14008 2021, CA Exec Order N-82-20 2020). While a majority of GCWA habitat conservation dollars have been spent conserving GCWA breeding habitat on the outskirts of the cities of Austin and San Antonio, our findings support previous work demonstrating higher rates of habitat conversion near metropolitan areas (Duarte et al. 2013). Given the scarcity of public lands, the distribution of intact forest habitat, and the relatively high amount of habitat loss and degradation occurring in and around metropolitan areas, future GCWA habitat conservation efforts should be more focused on supporting current protected areas and expanding protections to quality habitats in the Balcones Canyonlands and Fort Hood areas and regions west of San Antonio and Fort Worth. Additionally, projected species distribution models reflecting climate change impacts on tree species indicate that the Texas Hill Country will continue to be a stronghold for Ashe-juniper (with potential for population stabilization and maybe even increase/spread to the northeast; McKenney et al. 2007). This suggests that efforts to conserve or restore quality GCWA habitat will have long-term benefits.

We recognize the limitations of the analysis which equate all available mixed or evergreen forests within the breeding range, and not strictly those with Ashe-juniper components, as potentially suitable habitat for GCWA nesting. This is mainly due to current publicly available data sources and lack of LiDAR or other advanced geospatial datasets that would clarify spectral or structural differences in forest composition. Regardless, overall classification accuracies of the habitat loss dataset followed methods that average a mean absolute error of less than 3% (Kennedy et al. 2018). Additionally, datasets used for assessing habitat quality, though they represent the most current version available, are already out of date. Collectively, this indicates that our estimates for available habitat may be more liberal than in actuality. In the context of federal species listing and review, landscape change analyses, habitat identification and classification, and the characterization of trends over time must be considered. The metrics used in this study are meant to facilitate such consideration: they can be applied to multiple scales and interpreted by non-GIS audiences, helping diverse stakeholder groups to engage in the conservation decision making process. While current limitations in data, technology, and metrics may influence interpretation of landscapes, the intent is for future research to continue to improve upon this methodology and on our understanding of the changing habitat.

Human landscape modification is likely to continue in the Texas Hill Country, but conservation and land management actions can be taken to minimize further habitat loss and degradation in GCWA breeding range. This information will assist researchers and managers in prioritizing range-wide breeding habitat conservation efforts and highlights the significant role land management for conservation biodiversity plays on the landscape. There remains a need to grow the network of protected areas for GCWA restoration. Further, continued regular spatiotemporal assessments of habitat quantity and quality are necessary to assess changes to species potential for persistence and extrapolate population viability given these dynamics.

## ACKNOWLEDGEMENTS

Acknowledgements: We thank J Malcom, T Niederman and L O’Donnell for their thoughtful feedback in reviewing this work. We also thank the organizations that curate these publicly available datasets and the community scientists that work to collect these data. This research was conducted on secondary data sources. To our knowledge, no organisms were harmed in the collection of the datasets used.

